# Single-cell transcriptomics of malaria parasites

**DOI:** 10.1101/105015

**Authors:** Adam J. Reid, Arthur M. Talman, Hayley M. Bennett, Ana R. Gomes, Mandy J. Sanders, Christopher J. R. Illingworth, Oliver Billker, Matthew Berriman, Mara K. N. Lawniczak

## Abstract

Single-cell RNA-sequencing is revolutionising our understanding of seemingly homogeneous cell populations, but has not yet been applied to single cell organisms. Here, we established a method to successfully investigate transcriptional variation across individual malaria parasites. We discover an unexpected, discontinuous program of transcription during asexual growth previously masked by bulk analyses, and uncover novel variation among sexual stage parasites in their expression of gene families important in host-parasite interactions.

## Main text

Single-cell RNA-sequencing (scRNA-seq) has revealed astounding cell-to-cell heterogeneity, uncovered previously unknown cell types, and enhanced our understanding of developmental pathways in mammals ^1^. However, unicellular organisms have not yet been successfully explored using this method. Variation in the behaviour and developmental decision making of individual unicellular organisms cannot be resolved when averaging across populations of individuals. In *Plasmodium* parasites, transcriptomic analyses of isogenic strains grown under homogenous conditions revealed extensive transcriptional variation particularly among genes associated with host-parasite interactions ^2^. Furthermore, disease-relevant phenotypes such as commitment to sexual development ^3,4^, parasite sequestration ^5^ and nutrient acquisition ^6^ are thought to be driven by transcriptional variation between individual parasites.

Compared to mammalian cells, *Plasmodium* parasites present several challenges to existing single cell technologies, including much lower RNA content and a highly AT-biased base composition ^7^. Initially we trialled the well-established Smart-seq2 method ^8^ on sorted, *Plasmodium falciparum-*infected single red blood cells (Fig. 1a), but we increased the number of amplification cycles from 12 to 30 to account for their low RNA content. However, on average only 10% of reads mapped to genes in the parasite genome and more than half of these mapped to rRNA genes (Fig. 1b, left most bar). To improve yield, we used qPCR to evaluate the impact of modifying the anchor and length of the oligo(dT) primer, the reverse transcription enzyme, and the number of amplification cycles (Fig. 1c). A longer, unanchored polyT primer (T30) significantly improved yield (Fig. 1c), and was used in all further experiments. The reverse transcriptase enzymes SuperScript II and SMARTScribe gave the greatest yields by qPCR (Fig. 1c). Amplification by 25 and 30 cycles appeared equivalent by qPCR (Fig. 1c). Therefore we sequenced libraries generated from a small number of individual cells using the best two enzymes and either 25 or 30 cycles of PCR (Fig. 1b). The SMARTScribe enzyme with 25 or 30 cycles provided the best results, significantly increasing the number of genes detected and dramatically reducing the rRNA contamination (Fig. 1b; Supplementary Table 1). Given equivalent results for 25 or 30 cycles we opted to use fewer cycles for subsequent experiments. Transcript coverage was not affected by GC content or length; long genes were just as well detected as short genes, although we detected a slight 5’ bias in transcript coverage (Supplementary Figure 1). To validate our sorting protocol, a mixed population of *P. falciparum* and *P. berghei* schizonts were individually sorted into wells (n=72 cells). Reads from each parasite matched the expected species based on fluorescence suggesting that there was little cross cell contamination due to lysis (Fig. 1d; Supplementary Figure 2).

To begin to explore variation within parasite life cycle stages we generated 144 high quality single cell transcriptomes of mixed asexual and sexual (gametocyte) blood stage parasites of the rodent malaria model *P. berghei*. We identified expression from, on average, 1981 genes per cell, representing 33% of the total genes in the genome, on par with mammalian single cell experiments ^9^ (Supplementary Table 1; Supplementary Figure 3). Using a combination of Principal Components Analysis (PCA), k-means clustering ^10^ and comparison to bulk transcriptome datasets ^11,12^, we classified each cell as male, female, or asexual (Fig. 2a). The accuracy of our classification was strongly supported by established stage-specific markers (Fig. 2b). In addition to capturing sufficient transcriptional complexity to classify cells, we were also able to identify novel marker transcripts for each parasite stage (Supplementary Data 1).

**Figure 1.**
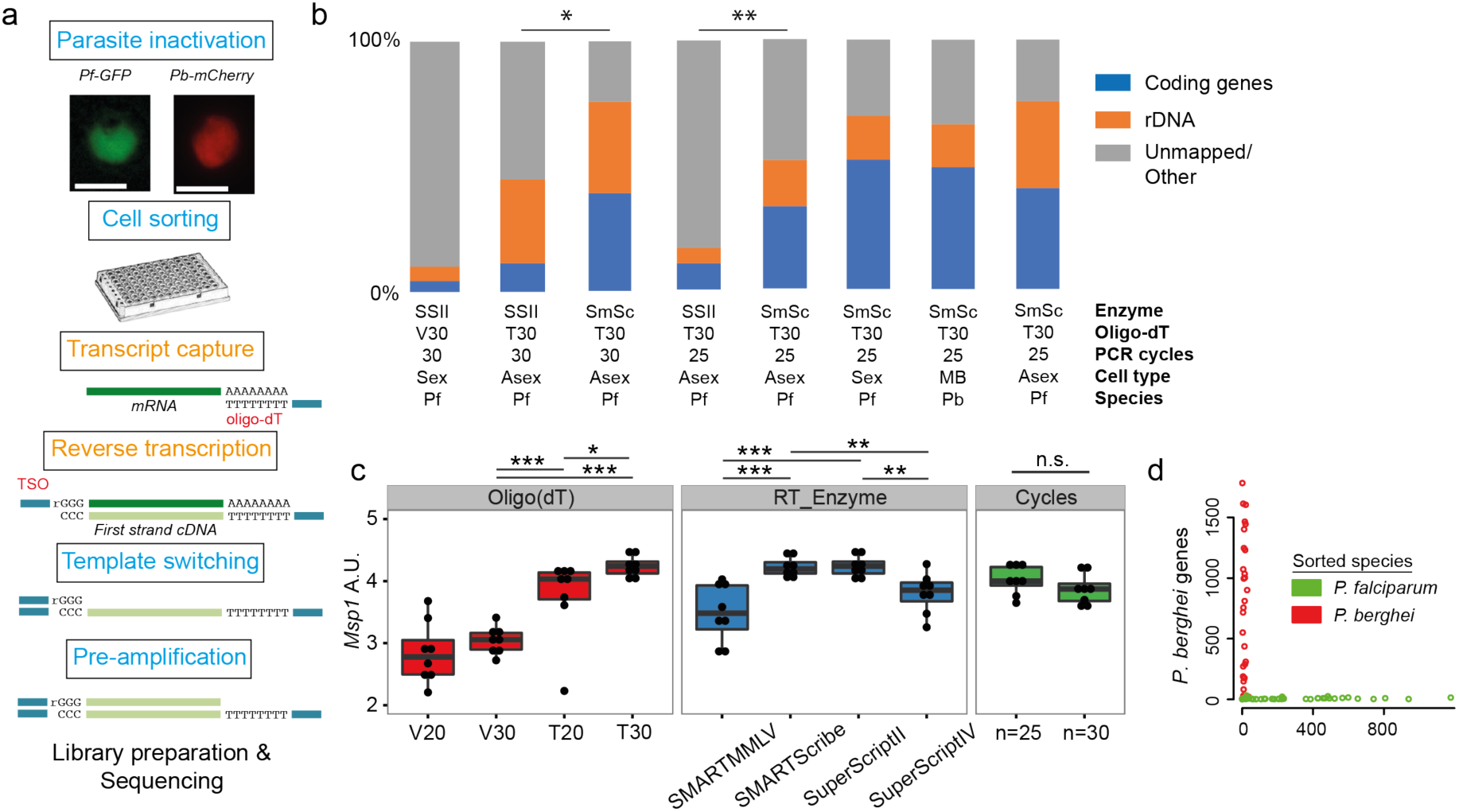
Establishment of a robust protocol for single cell transcriptomic analysis of Plasmodium parasites. ***(a)** Overview of the single cell RNAseq protocol. Steps in the original Smart-seq2 protocol ^8^ that resulted in significant gains are highlighted in orange. (**b**) Relative numbers of reads mapping to coding RNA and rDNA for each trial, averaged over all cells in that trial. The final three bars represent the main P. falciparum gametocyte, P. berghei mixed blood and P. falciparum asexual datasets, respectively. Asterisks indicate selected significant differences between proportions of reads mapping to coding genes, calculated using Mann-Whitney U. (**c**) The protocol was evaluated using qPCR of the msp-1 transcript (PF3D7_09303000) on sorted pools of 10 asexual parasites (n=8) (Significance from Mann Whitney test). The following reagents were tested: Oligo(dT)s containing a terminal anchoring base (A,G,C; V) or not (T) and of varying lengths (20 Ts vs. 30 Ts); 4 reverse transcriptase enzymes; 25 or 30 cycles of preamplification. (d) Individually sorted P. falciparum and P. berghei cells from a mixed pool revealed no doublets and little contamination (see Supplementary Figure 2).*

Whereas mature sexual parasite cells are terminally differentiated, transcriptional variation along the 24 hour asexual cycle of *P. berghei* is thought to be continuous ^11,13^, including during the transition from the growth phase (trophozoite) to the budding phase (schizont).

We predicted the maturation state of each individual asexual cell, using a rank-correlation approach to compare them to two published bulk RNA-seq datasets ^11,12^. We also carried out a pseudotime analysis ^14^ using variable genes ^15^ to order the asexual cell subset. We observed a strong concordance between the temporal and pseudotemporal predictions (Fig. 2c and 2d). Interestingly, we observed very abrupt transcriptional changes, distinct from the smooth transitions observed in bulk time course experiments (Fig. 2d, Supplementary Figure 4). To further study the dynamics of transcriptional shifts over asexual development, we analysed the transcriptomes of 155 *P. falciparum* late asexual stages. These cells could also be ordered in pseudotime in agreement with their predicted stage (Fig. 2e; Supplementary Figure 5) and displayed the same dramatic transcriptional profile shift in the late asexual cycle (Supplementary Figure 5). The smooth transitions observed in bulk data may be attributable to averaging across slightly different asexual life cycle points, which does not happen when examining single cells in pseudotime. Taking advantage of our high resolution transcriptomes, we conducted a co-expression correlation analysis of the ApiAp2 family of transcription factors (TFs) ^16^ and observed marked correlated and anticorrelated clusters in a TF network (Supplementary Figure 6). These patterns were strongly associated with peak expression along pseudotime and several correlations were conserved in both species (Supplementary Figure 6). This suggests a possible conserved regulatory framework underlying this rapid and discrete transcriptional shift in late schizogony. More comprehensive sampling of the asexual cycle will be necessary to fully evaluate the pace of transitions and the role of synergistic or antagonistic interactions between TFs in establishing the discrete patterns of gene regulation we observe during the asexual cycle of *Plasmodium* parasites.

**Figure 2.**
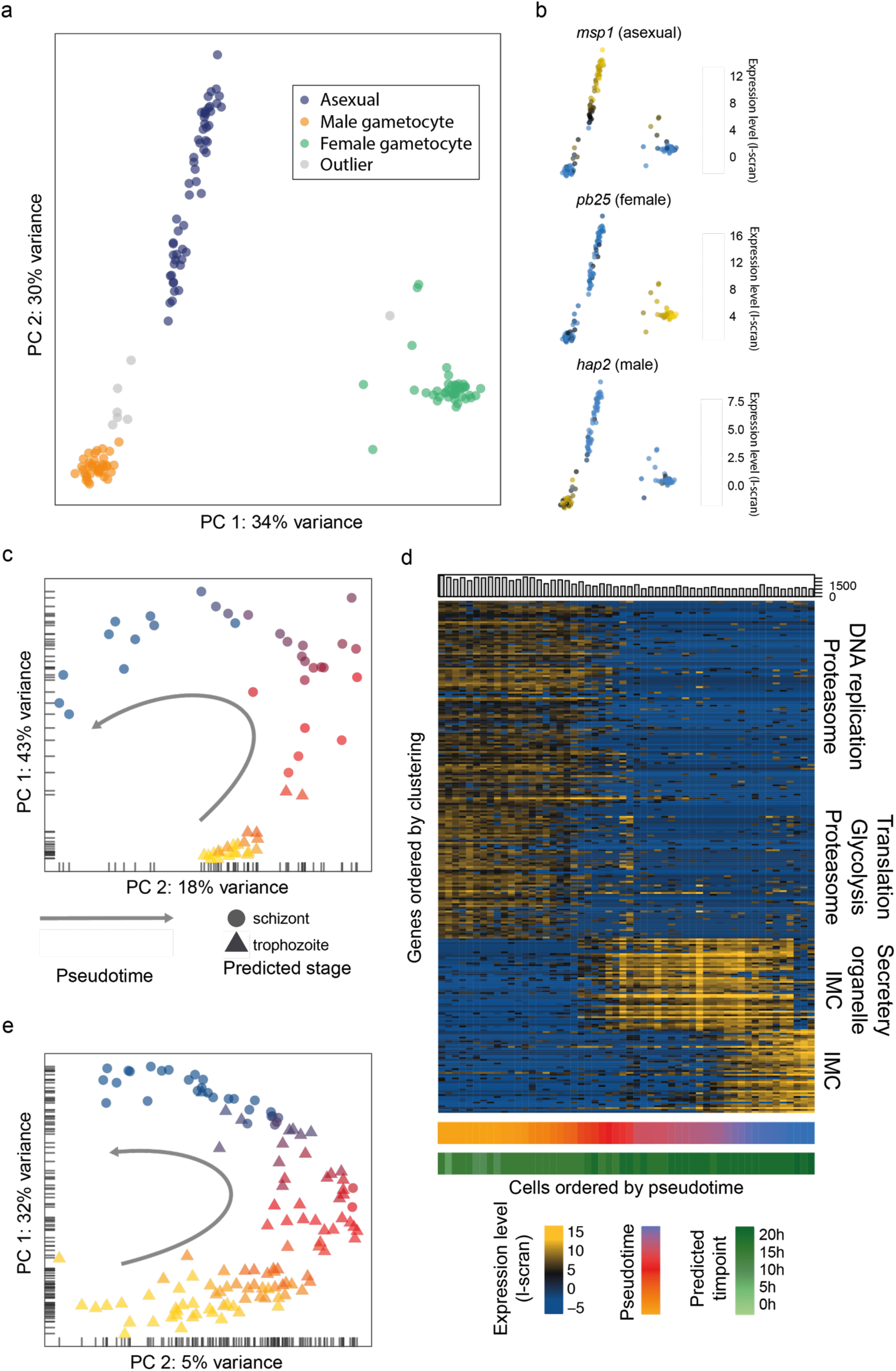
Single cell RNA-seq allows dissection of parasite populations. ***(a)** A combination of Principal Components Analysis (PCA), k-means clustering and comparison to bulk RNA-seq datasets was used to classify 144 high quality P. berghei single cells, and revealed three distinct subpopulations. Outliers may represent erythrocytes infected with both sexual and asexual stages or early stages in gametocyte development. **(b)** Three well-established markers of the male, female and asexual lineages ^21-23^ are concordant with our classification. **(c)** Pseudotime ordering (using ^14^) of the asexual cells in (a) was in close agreement with bulk RNA-seq datasets (predicted timepoint from ^11^, predicted stage = consensus; see Methods). **(d)** Differentially expressed genes (identified using ^15^) were clustered along pseudotime revealing groups of genes with abrupt expression profile changes during late asexual cycle. Functional enrichment in the clusters was in agreement with the expected shift from the growing trophozoite to the budding schizont (IMC = Inner Membrane Complex). **(e)** Pseudotime ordering (using ^14^) of the 125 P. falciparum late asexual cells was in close agreement with bulk RNA-seq datasets (predicted timepoint from ^24^, predicted stage = consensus; see Methods).*

Transcriptional variation within large gene families is known to be important for immune evasion and establishment of chronic infection by *Plasmodium* in the mammalian host ^17,18^. However, relatively little is known about the role of these families, or host-parasite interactions more generally, in the sexual stages which are transmitted to the mosquito. We examined transcriptional variation within the four *P. berghei* life stages and found 115 variable genes in females, 73 in males, 27 in trophozoites and 4 in schizonts (Supplementary Data 2). A full functional description of all variable genes is described in Supplementary Data 3. In trophozoites, we observed variation in expression of the multigene family *pir*, in particular the L-type *pir* genes ^18^. Variation in expression of these genes has recently been associated with establishment of chronic infection ^18^. In male gametocytes, but not females, we observed variability of S-type *pir* genes ^12^, a group for which no function has yet been described (Fig. 3a; Supplementary Data 4). Their products however, have been identified in male gametes ^19^, the cells that arise from male gametocytes once ingested by a mosquito. Our data suggest that sex-specific variation in expression of these genes may play a role in malaria parasite transmission.

**Figure 3.**
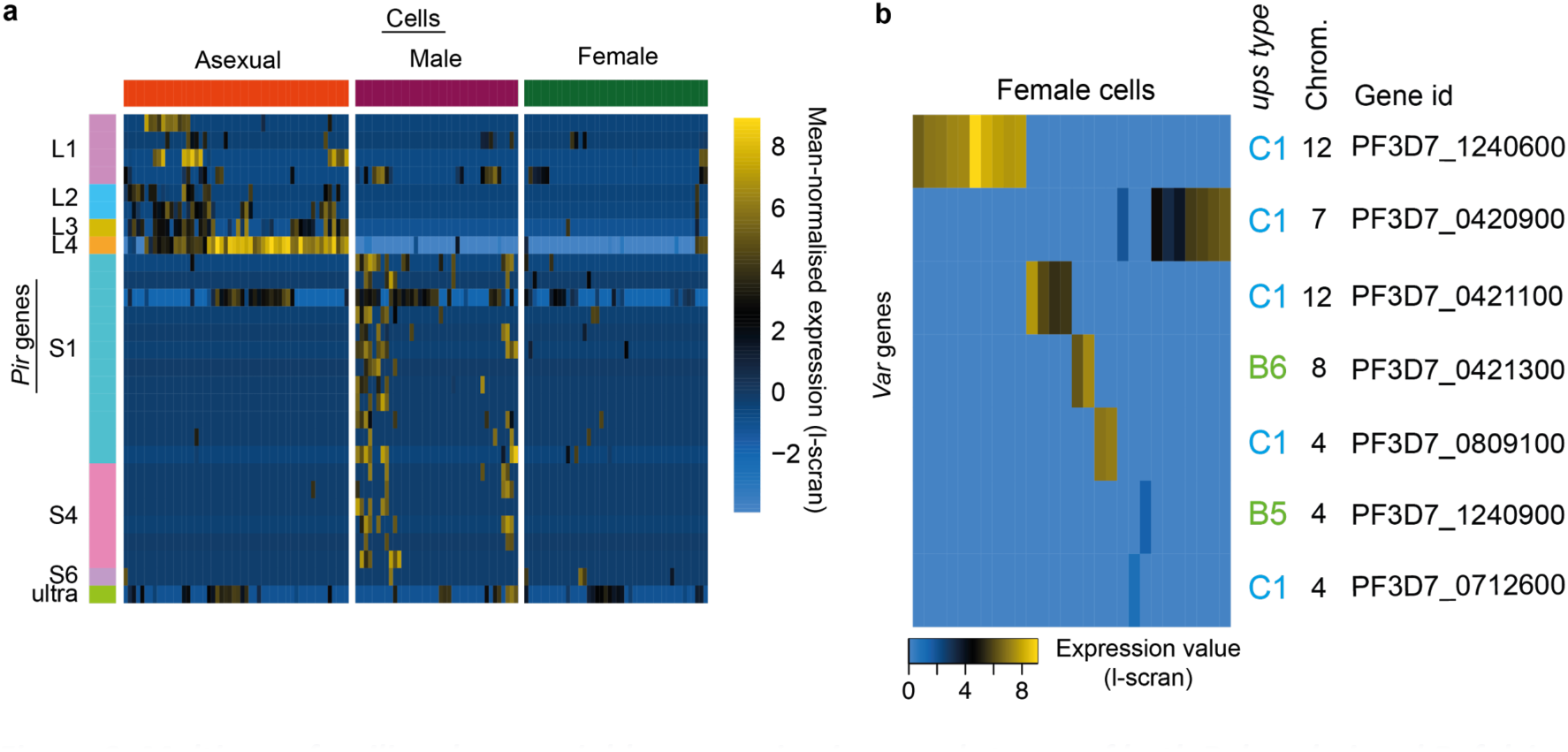
Multigene families show variable expression in sexual stages of both P. berghei and P. falciparum. **(a)** Pir gene expression was highly variable across male gametocytes. In addition, more pir genes were expressed in males than females. These are distinct subfamilies of pir genes from those variably expressed in asexual stages. **(b)** 28 female P. falciparum gametocytes expressed var gene transcripts from the sense strand. We only ever detected a single mRNA var in each cell, suggesting the maintenance of mutually exclusive var expression in gametocytes. The promoters of expressed var genes were only upsC and upsB subtypes from internal rather than subtelomeric regions of the genome.

To further explore transcriptional variation in sexual stage biology, we carried out scRNA-seq on *P. falciparum* late sexual stage individuals, resulting in 191 high quality single cell transcriptomes (Supplementary Table 1, Supplementary Figure 3). Due to the small number of predicted males detected (n=5; Supplementary Figure 7), we only examined expression variation in females, detecting 448 variable genes (Supplementary Data 2). The repertoire of multigene families in *P. falciparum* is highly diverged from *P. berghei*^20^ and clear *pir* gene orthologues do not exist. However, we found an enrichment for variation in the well-studied *var* gene family (14 of 60 genes, hypergeometric test, *p* = 0.0006). Expression of noncoding *var* transcripts is common and is involved in maintaining mutually exclusive expression of a single *var* gene ^17^. Therefore, to assess the presence of true *var* mRNAs, we identified reads that confirmed intronic splicing. We discovered that although there were reads mapping to multiple *vars* within each cell, only a single *var* gene ever had reads that spanned the spliced intron. This suggests that mutually exclusive expression of *var* genes occurs in sexual stages, as it does for asexual parasites ^17^. The *var* genes variably expressed across females were always from internal *var* gene clusters, consisting primarily of *upsC* type *vars* (Fig. 3b) but whose role is largely unexplored. In asexual stages, *var* gene products (PfEMP1) allow the parasites to adhere to the host vasculature and avoid clearance in the spleen and are important in evading the adaptive immune system ^17^. Variable and mutually exclusive transcription of *upsC-rich* internal *var* genes in female gametocytes suggest that this variable gene family, like the *pir* genes above, may also play an important role in parasite transmission.

In this study, we have demonstrated a highly successful approach to single cell RNA-sequencing in the study of a unicellular eukaryotic organism. We have used scRNA-seq to successfully resolve individual parasites within a mixed population, to classify them, and to identify previously hidden transcriptional variation. Our data suggest that the transcriptional cascade during development in red blood cells is not continuous as previously described, but instead it occurs in discrete stages. This has important implications for our understanding of gene regulation in the most pathogenic stage of the parasite. We also show that variable expression of genes involved in host-parasite interactions is a feature of multiple life stages of *Plasmodium* parasites. We hope that our optimisation framework will assist in extending scRNA-seq to a much wider range of diverse eukaryotic cell types.

## Methods

### Isolation of *P. berghei* parasites

The constitutively mCherry-expressing *P. berghei* ANKA line, clone RMgm-928 ^25^, was propagated in a female 6- to 8-week-old Theiler’s original outbred mouse supplied by Envigo UK. Parasites were purified from an overnight (20h) 50 mL culture of 1 mL of infected blood using a 55% Histodenz cushion (SIGMA), following an established schizont purification protocol detailed elsewhere ^26^. Purified late stages (asexual and sexual) were pelleted at 450g for 3 minutes and incubated with 500 µL of RNALater (ThermoFisher) for 5 minutes, and further diluted into 3 mL of 1x PBS prior to cell sorting. All animal research was conducted under licenses from the UK Home Office and used protocols approved by the ethics committee of the Wellcome Trust Sanger Institute.

### *In vitro P. falciparum* culture

3D7-HTGFP, a GFP-expressing *P. falciparum* strain ^27^, was maintained in O-negative red blood cells obtained from the NHSBT, using RPMI 1640 culture medium (GIBCO) supplemented with 25 mM HEPES (SIGMA), 10mM D-Glucose (SIGMA), 50 mg/L hypoxanthine (SIGMA), 10% human serum (obtained locally in accordance with ethically approved protocols), and gassed using a mix containing 5% O^2^, 5% CO^2^ and 90% N_2_. Parasites were highly synchronised using two consecutive cycles of Percoll-Sorbitol treatment ^28^. Late asexual parasites (trophozoites and schizonts) were purified on a cushion of 63% Percoll (GE Healthcare). Stage V gametocytes were obtained using standard gametocyte culturing ^29^ and purified magnetically with an LS column ^30^ (Miltenyi Biotec). Following purification of each stage, all *P. falciparum* parasites were pelleted at 800g for 5 minutes, incubated with 500 µL of RNALater (ThermoFisher) for 5 minutes, and further diluted into 3 mL of 1x PBS prior to cell sorting. Parasitaemia was determined by Giemsa–stained thin blood smear.

### Cell sorting

4 µl of lysis buffer (0.8 % of RNAse-free Triton-X (Fisher) in nuclease-free water (Ambion)), UV-treated for 30 min with a Stratalinker UV Crosslinker 2400 at 200, 000 µJ/cm2, 2.5 mM dNTPs (Life Technologies), 2.5 µM of Oligo(dT) (IDT; see Supplementary Table 1 for detail) and 2U of SuperRNAsin (Life Technologies) were dispensed into each well of the recipient RNAse-free 96-well plate (Abgene) immediately prior to the sort and kept on ice. In the first experiment only 2 µl of lysis buffer were used but the observed cell-capture efficiency was very poor so the volume was increased. Cell sorting was conducted on an Influx cell sorter (BD Biosciences) with a 70 µm nozzle. Parasites were sorted by gating on single cell events and on GFP (*P. falciparum*) or mCherry (*P. berghei*) fluorescence. A non-sorted negative control well and a positive 100-cell control well were included in every plate alongside single cells. Sorted plates were spun at 200 G for 10 seconds and immediately placed on dry ice.

### First and second strand cDNA synthesis and pre-amplification

Cells in plates were incubated at 72 °C for 3 minutes. A reverse transcription master mix was added to the samples containing 1 µM of LNA-oligonucleotide (5’-AGCAGTGGTATCAACGCAGAGTACATrGrG+G-3’; Exiqon), 6 µM MgCl_2_, 1M Betaine (Affymetrix), 1× reverse transcription buffer, 50 µM DTT, 0.5 U of SuperRNAsin, and 0.5 µl of reverse transcriptase (Supplementary Table 1). The total volume of the reaction was 10µl. The plate was incubated using the following programme 1 х 42°C/90’; 10 х (42°C/2’, 50°C/2’), 1 х 70°C/15’. Samples were then supplemented with 1х KAPA Hotstart HiFiReadymix and 2.5 µM of the ISO SMART primer ^8^ and incubated using the following cycling programme 1 х 98°C/ 3’; 25 or 30 х (98°C/20’’, 67°C/15’’, 72°C/6’); 1х 72°C/ 5’ (Supplementary Table 1). Samples were then purified with 1х Agencourt Ampure beads (Beckman Coulter) in a Zephyr G3 SPE Workstation (Perkin Elmer) according to the manufacturer’s recommendation. Amplified cDNA was eluted in 10 µl nuclease-free water. Details of different permutations of the protocol tested during the optimisation process are given in Supplementary Table 1.

### Quality control of cDNA samples

The quality of a subset of amplified cDNA samples was monitored with the high-sensitivity DNA chip on an Agilent 2100 Bioanalyser. Samples were verified by qPCR using LightCycler 480 SYBR Green I Master and MSP-1 primers at a concentration of 0.4 µM (Forward: 5’-TCCCAATCAGGAGAAACAGAAG-3’; Reverse: 5’-GATGGTTGTGTTGGTGGTAATG-3’), on a Roche Lightcycler 480 II. Reactions were incubated according to the following cycling programme: 1x 95°C/10’; 45X (98°C/20’’, 58°C/10’’, 68°C/30’’). Transcripts were quantified with the absolute quantification method using a standard dilution.

### Library preparation and sequencing

Libraries were prepared using the Nextera XT kit (Illumina) according to manufacturer recommendations. 96 or 384 different index combinations were used to allow multiplexing during sequencing. After indexing, libraries were pooled for clean-up at a 4:5 ratio of Agencourt Ampure beads (Beckman Coulter). Quality of the libraries was monitored with the high sensitivity DNA chip on an Agilent 2100 Bioanalyser. Empty-well controls and single cells were pooled separately from 100 cell controls and loaded proportionally to their expected cell content for sequencing on an Illumina MiSeq or HISeq 2500.

### Sequencing of single-cell libraries

The original Smart-seq2 protocol with the Superscript II enzyme and the original Oligo(dT) with an anchoring base was run with 30 PCR cycles of preamplification on 10 samples. The samples included a single no-cell control, five single *P. falciparum* gametocytes, two 10-cell controls and two 100-cell controls. These were multiplexed, along with three samples each of individual human lung carcinoma cells (A549) and sequenced on a single MiSeq run with 150bp paired end reads.

To test the effect of different reverse transcriptase enzymes and different numbers of PCR cycles, we sequenced *P. falciparum* schizont libraries prepared using 1) the SmartScribe enzyme (Clontech) for a) six single cells, one 100-cell control and two no-cell controls with 25 cycles of PCR, and b) six single cells and one 100-cell control with 30 cycles of PCR; and 2) the SuperScript II enzyme (Thermofisher) for a) six single cells, one 100-cell control and two no-cell controls with 25 cycles of PCR, and b) six single cells and one 100-cell control with 30 cycles of PCR. These were multiplexed on a single MiSeq run with 150bp paired end reads.

To determine whether single-cell samples might be contaminated with either additional cells or RNA from lysed cells, individual mCherry *P. berghei* (RMgm-928 ^25^) and GFP *P. falciparum*^*27*^ schizonts were mixed in a 1:1 ratio, inactivated with RNALater fixation and then sorted. A multiplex library was prepared comprising 32 single *P. berghei* schizonts, two 100-cell *P. berghei* schizont controls, one no-cell control, 40 single *P. falciparum* schizonts and two 100-cell *P. falciparum* schizont controls. These libraries were sequenced as a multiplex pool on a single MiSeq run with 150bp paired end reads.

The *P. berghei* mixed blood stage samples comprised 182 single-cells of *P. berghei*, plus four no-cell controls and six 100-cell controls. These were multiplexed with another 192 samples not analysed in this work and sequenced on a single HiSeq 2500 lane using HiSeq v4 with 75bp paired end reads. The *P. falciparum* gametocyte samples were sequenced as three multiplexed pools of 84, using the same chemistry. Three technical duplicate samples were excluded from analysis. The *P. falciparum* asexual samples were sequenced as two pools of 96, each on one Illumina HiSeq 2500 lane using HiSeq v4 chemistry with 75bp paired-end reads. Each batch of 96 samples contained three 100-cell controls. The second batch (lane 7) contained six samples of stage I gametocytes and six samples of stage II gametocytes, each with a single 100-cell control. These were not included in the analysis, leaving 176 single cell samples.

### Mapping reads and calculating read counts

All sequencing experiments were processed in the following way. CRAM files of reads were acquired from the WTSI core pipeline, converted to BAM using *samtools-1.2 view -b*, sorted using *samtools sort –n*, converted to fastq using *samtools-1.2 bam2fq* and then deinterleaved ^31^. Nextera adaptor sequences were trimmed using *trim_galore -q 20 -a CTGTCTCTTATACACATCT ––paired ––stringency 3––length 50-e 0.1* (v0.4.1). HISAT2 (v2.0.0-beta)^32^ indexes were produced for the *P. falciparum* v3 (http://www.genedb.org/Homepage/Pfalciparum) or *P. berghei* v3 ^33^ genome sequences, downloaded from GeneDB ^34^, using default parameters (October 2016). Trimmed, paired reads were mapped to either genome sequence using *hisat2 ––max-intronlen 5000 -p 12*. For the dual sort experiment we mapped against a combined reference, allowing us to exclude reads that map to both genomes. SAM files were converted to BAM using *samtools-1.2 view –b* and sorted with *samtools-1.2 sort*. GFF files were downloaded from GeneDB (October 2016) and converted to GTF files using an in-house script. All feature types (mRNA, rRNA, tRNA, snRNA, snoRNA, pseudogenic_transcript and ncRNA) were conserved, with their individual “coding” regions labelled as CDS in every case for convenience. Where multiple transcripts were annotated for an individual gene, only the primary transcript was considered. Reads were summed against genes using HTSeq: *htseq-count-f bam-r pos-s no-t CDS* (v0.6.0;^35^). Multimapping reads were excluded by default (-a 10). For downstream analysis (excluding examination of rRNA counts) transcripts not included in the GeneDB cDNA sequence files were excluded. The read counts for *P. berghei* mixed blood stages, *P. falciparum* gametocytes and *P. falciparum* asexual stages are presented in Supplementary Data 6.

### Classifying reads for quality control

To determine the useful yield of different RNA amplification protocols (summarized in Supplementary Table 1), we classified resulting reads into those mapping to rRNA genes, other genes, unmapped or ambiguous (falling into more than one category). We concentrated here on rRNA because we had observed that this was a particular problem. To do this we began with HISAT2 BAM files produced as described above. Total read pairs were all the unique read pair ids. Ribosomal RNA reads were counted using *bedtools intersect* (v2.17.0; ^36^) to find the overlap of unique read pair ids with rRNA features. Other coding reads were counted in the same way, but looking for overlap with all other features. Unmapped reads were identified using *samtools view -f 0х8* (v1.2) and extracting unique read pair ids. Where a read pair occurred in more than one of these lists, it was counted as ambiguous.

We compared the library complexity of different iterations of our protocol in order to determine whether more reads resulted in more complexity, or simply more reads from the same genes, perhaps due to large numbers of PCR cycles. Different sequencing runs had very different library sizes and so we downsampled the data. To maximise the number of cells included, while also allowing a reasonable number of reads per cell, we chose to downsample to 50000 reads per cell. To do this, 50000 counts from HTSeq were randomly sampled for each cell. Counts associated with protein coding genes were enumerated and genes were called as detected if there were at least 10 reads mapping to them.

### Assessing bias in single-cell sequencing libraries

Different library preparation and sequencing protocols exhibit different biases in representation of GC/AT-rich sequences and 5’ or 3’ transcript ends. In order to assess such biases we took an approach of using the mapped RNA-seq data to identify fragments of genes which were expressed and examined the coverage of genes by these fragments. The reason for doing this, rather than looking at coverage depth was that we had noticed that genes often did not have full coverage, particularly when very long or expressed at a low level. This suggests that, although we would expect Smartseq-2 to amplify full length transcripts, in some cases only partial transcripts survived the full protocol. We used Stringtie (v1.2; default options;^37^) to call expressed fragments from our HISAT2 BAM files. We then looked for Stringtie transcript features overlapping each mRNA feature in our reference annotation. Where multiple Stringtie transcripts overlapped each other, these were merged. We then determined, for each gene, the exonic sequence covered by the merged Stringtie transcripts. The length, GC content and relative start and end of these regions was calculated. Observed GC content was compared against the GC content for the whole coding region. Each relative position along a coding sequence (0-100) covered by a fragment was incremented for each fragment covering it. The coverage of each relative position for each gene was then normalised between 0 and 1 based on the highest coverage across that coding sequence. To examine the effect of gene length, we compared the length distribution of all 4943 *P. berghei* genes used in our initial analysis to the 4579 which passed our filtering criteria (having at least 10 reads in at least 5 cells).

### Analysis of contamination with a dual sort of *P. falciparum* and *P. berghei* schizonts

Reads for the dual sort samples were mapped as above, but to a combined reference of both parasites, enabling reads that map equally well to both genomes to be discarded as their origin could not be determined. Read counts were converted to FPKMs and transcripts with an FPKM >= 10 were counted as expressed. We used these data to show that no well contained more than one cell, i.e. wells with good data (a large number of expressed genes) never had similar numbers of genes from both species. Furthermore no good wells contained a large number of genes from the incorrect species. To explore whether contaminating genes were similar in different wells, we compared *P. falciparum* genes identified in wells with a *P. berghei* cell sorted into them and vice versa between wells. Similarity was calculated as the number of common contaminating genes with an FPKM >= 10, divided by the average number of contaminating genes between the two wells. Each cell contained relatively few contaminating genes. This was higher for *P. falciparum* contamination of *P. berghei* (Supplementary Figure 2b) than *P. berghei* contamination of *P. falciparum* (Supplementary Figure 2c), suggesting *P. falciparum* cells contribute more to extracellular RNA in the medium. Different cells shared relatively few contaminating transcripts, but the more commonly occurring contaminants were also more highly expressed in their cells of origin (Supplementary Figures 2d,e). The contamination in cells was generally very low and reflected the amount of RNA present in the cells of origin.

### Filtering and normalisation of single-cell read count data

The three main datasets (Pb mixed, Pf asex, Pf sex) were processed using Scater v1.0.4 ^38^. Firstly we removed genes with no counts in any cell. We then removed remaining control cells, cells with a total of less than or equal to 25000 read counts and/or less than 1000 genes with at least one read. Subsequently we removed genes which did not have at least 10 reads in 5 cells. For the *P. berghei* dataset this resulted in 144/183 cells and 4579 unique genes detected across all cells. For the *P. falciparum* gametocyte dataset there were 191/238 cells and 4454 unique genes after filtering and for the *P. falciparum* asexual dataset 161/180 cells and 4387 unique genes. The counts were then normalised using scran ^39^ (v1.0.3). Normalisation is required due to technical variation between samples due to, for example, variable sequencing depth and capture efficiency. Single cell RNA-seq read count data contain many zeroes compared to bulk RNA-seq data. These are caused by drop out of low expressed genes or variation between cells and reduce the accuracy of normalisation methods designed for bulk RNA-seq data. Scran uses a pooling approach to reduce these zeroes. Furthermore, it allows an initial clustering of the data and normalisation within these clusters (e.g. cell types), prior to a final normalisation step across the whole dataset. This is particularly useful for our *P. berghei* data, where the asexual, male and female gametocyte cells differ greatly in their expression patterns. The initial clustering step was performed with the *scran* function *quickCluster* (min.size = 30). This resulted in three clusters representing the asexual, male and female gametocyte populations. The *computeSumFactors* function was run using these clusters, with sizes = 20 and positive = TRUE. All downstream analyses were performed with the scran normalised data except where stated. For *P. falciparum* gametocytes, the *computeSumFactors* function was run with sizes = 15. For *P. falciparum* asexual stages, we set min.size = 20 for quickCluster and the *computeSumFactors* function was run with sizes = 10.

For some applications it is necessary to normalise the data by transcript length. For instance, when comparing ranked gene expression values to reference data for determining life cycle stage of a cell. We therefore normalised the scran values by taking the exponent (2^x^), multiplying by 1000 and dividing by the cDNA length, determined from the GeneDB cDNA FASTA file (coding sequence only, no UTRs). This is similar to the FPKM calculation, except the library size normalisation is already accomplished. We refer to these values as *l-scran*, for length-normalised scran values.

### Determining parasite life cycle stages using bulk reference data and clustering

We used several bulk RNA-seq data sets to assign a life cycle stage to each cell. For *P. berghei* asexual stages we used microarray data ^11^ that captures the 24-hour asexual development cycle at two-hour resolution. In their experiment Cy5 was used to label each time point while Cy3 was used to label a pool of all samples. The “F635 Median-B635” values are the difference in Cy5 intensity between the median foreground and the median background. This intensity value is related to the actual expression level and these are the values we used. Their data were generated using the *P. berghei* v2 genome assembly, so we remapped their probe sequences against v3 using HISAT2 (default parameters). We then used *htseq-count -a 200 -f sam -r name -s no* to identify the genes to which the probes mapped (cut -f1,21 probes_berghei_htseq.sam | grep PBANKA | grep -v ambiguous > probes_berghei.map). We then used the GPR files provided from ArrayExpress ^40^ (accession GSE80015) and the probe map to produce a table of percentile ranks for each gene in each condition.

Single cell gene expression values were converted to length normalised scran values (l-scran), as described above, in order to produce more accurate rank expression levels for our scRNA-seq data. We compared each single-cell expression profile against each reference data set. To reduce noise, genes that do not vary greatly between conditions in the reference data were removed. For the *P. berghei* 24h intraerythrocytic developmental cycle reference data ^11^, genes were only included if their expression profile had a mean rank of greater than 30 and less than 70 and standard deviation in rank across samples of greater than 3. Genes from the query dataset with l-scran < 3 were also removed. A minimum of 100 remaining genes common to both the reference and query profiles were required to calculate a correlation between them. The Spearman rank correlation was used in order make the microarray and RNA-seq datasets more comparable. The best correlation of a single-cell expression profile with a reference expression profile was taken as the consensus stage prediction for that single-cell. As new data (e.g. single-cell analysis of timepoints across the full, synchronised erythrocytic development cycle) become available, benchmarking staging algorithms will become feasible. Bulk RNA-seq data to classify *P. berghei* males and females directly was not available. Therefore, we used bulk RNA-seq data ^12^ that includes mixed-sex gametocyte samples, after converting the profiles to v3 using previous id annotation from PlasmoDB ^41^.

To determine distinct groups of single-cells based on their expression patterns we used the clustering tool SC3 ^10^. We used the combined Euclidean, Pearson and Spearman distance, plus the combined PCA and spectral transformation. For the *P. berghei* dataset the optimal *k* was 3 (average silhouette width = 0.99), with 4 being nearly as good (average silhouette width = 0.97). We found that the additional cluster split the asexual parasites into trophozoites and schizonts, while both *k* values retained the male and female gametocytes as separate clusters. However, there was still extensive variation within these clusters so we further investigated this by excluding asexual cells and clustering again. With this reduced dataset we were able to get a new, robust clustering with k=3 (width = 0.99). Here, outliers from both the putative male and female clusters clustered together, exclusive of the core of male and female clusters. Markers suggested that six of these outlier cells possessed both male genes and asexual genes, while a single cell possessed both female genes and asexual genes. It is possible that these cells are early gametocytes, committed schizonts or cells doubly infected with both asexual and sexual parasites. These were excluded from further analysis. The *markers* function of SC3 (AUROC threshold 0.85, p-value threshold 0.01) was used on the initial clustering, with k = 3, to identify novel markers for asexuals, males and females.

Data from Lasonder and colleagues ^42^ was used to classify *P. falciparum* gametocyte cells by sex. Raw count data was downloaded from the Gene Expression Omnibus ^43^(accession GSE75795) and converted to FPKM. Data from Young and colleagues ^44^, was used to classify *P. falciparum* cells along the gametocyte development time course (days 1, 2, 3, 6, 8, 12). For this dataset profiles of ranks were downloaded from PlasmoDB. The Lasonder data ^42^ highlighted five male cells, with the rest called as females. The Young data ^44^ suggested that all the cells were at a consistent stage of development (eight days), although resolution is lacking at the most relevant timepoints, between eight and twelve days. The classification of each cell is listed in Supplementary Data 5.

Bulk RNA-seq data from Otto *et al.* ^24^ and Lopez-Barragan *et al.* ^45^ were used to classify 161 *P. falciparum* asexual stage cells. RNA-seq reads from Otto et al. ^24^ for the 36bp Illumina libraries only, were downloaded from the European Nucleotide Archive (accession ERX001048). They were mapped to the *P. falciparum* 3D7 genome sequence using HISAT2 v2.0.0-beta ^32^ and reads were counted using HTSeq v0.6.0 ^35^. Read counts were then converted into FPKM for subsequent analysis. RNA-seq reads from Lopez-Barragan *et al.* ^45^ were downloaded from the European Nucleotide Archive (accession SRX105940) and mapped to *P. falciparum* 3D7 transcript sequences using Bowtie2 v2.2.9 (-a -X 800; ^46^) and eXpress v1.5.1 ^47^. The resulting read counts were converted to FPKM. The Lopez-Barragan prediction ^45^ was used as the consensus prediction, the prediction included 6 stage II gametocytes which were removed from further pseudotime analysis (n=155).

### Assessment of gene expression variation during asexual maturation

Within the 54 *P. berghei* cells identified as asexual, 277 genes were found to be variable using M3Drop ^15^ (raw count input, False Discovery Rate <= 0.01). L-scran expression values for this subset of genes (expressionFamily=tobit()) was used to order the cells in pseudotime using Monocle 2 ^14^; specifically the reduceDimension() and orderCells (num_path=2) functions were used to derive the ordering of the cells. Monocole 2 identified a single cell state and the cells were ordered in a single trajectory (Fig. 2c). The Monocle 2 package was further used to cluster genes in pseudotime (k-means) with the clusterGenes() function. We looked for enrichment of Gene Ontology terms within the four clusters identified, using topGO ^48^ (summarized in Fig. 2d).

For the *P. falciparum* 155 cell dataset, 361 genes were found to be variable genes with M3Drop (raw count input, False Discovery Rate <= 0.01) ^15^, Monocle 2 identified 2 branches defining three possible trajectories, although 2 of those appeared minor (Supplementary Figure 5). Cells ordered in these minor trajectories did not seem to correlate with known biological markers, such as sexual commitment markers (*ap2-g* and *gvd-1*; Supplementary Figure 5) and were removed for the rest of the analysis. The pseudotime analysis was repeated on the main trajectory cells (125 cells) (Supplementary Figure 5). The Monocle 2 package was further used to cluster genes in pseudotime (k-means) with the clusterGenes() function.

### Direct comparison of single-cells in pseudotime with bulk RNA-seq data

To determine whether the same set of genes displayed different patterns across development in bulk and single-cell RNA-seq experiments we made direct comparisons between these two approaches. After ordering the *P. berghei* asexual cells by pseudotime, genes were ordered by their peak of expression based on linear (i.e. not logged), length-normalised scran expression values. To do this, expression value data, ordered by pseudotime, were normalised, then Fourier transformed, sorting transcripts according to the phase of the most prominent frequency. Signal-to-noise (S/N) ratios were calculated for each transformed signal and normalised with respect to the maximum achievable value for the dataset. Transforms with a normalised S/N of less than 0.1 were excluded from the results as lacking evidence of periodicity. The Hoo *et al.* dataset ^11^ were treated in the same way, but initially ordered by time point of collection rather than pseudotime and using intensity values as described above. This resulted in 1141 ordered genes for our single cell data and 2612 genes for the Hoo data ^11^. There were 651 shared genes, which were used to compare the two datasets.

The *P. falciparum* asexual cells were ordered differently. We used the Otto *P. falciparum* asexual development cycle time course data ^24^ as a reference. These data were processed as described above. We used the Fourier transform approach described above, with a normalised S/N ratio of 0.5 to identify 4517 genes from the Otto et al. dataset ^24^. We then identified 336 genes common to this list and the list of 361 differentially expressed genes identified across the 155 single cells. This approach was taken because the window of time captured by our single-cells was too narrow to identify cycling genes using the Fourier approach (all the normalised S/N ratios were very low e.g. <0.05). We then generated heatmaps for the two datasets, with the genes ordered by their peak time in the Otto et al. dataset ^24^ in both cases.

### Determining gene expression variability within different cell types

To examine gene expression variation within life stages, we used the filtered datasets and considered male, female, trophozoite and schizont cells separately. For *P. berghei*, M3Drop ^15^ was used to determine gene expression variability amongst cells with FDR <= 0.05. We identified 115 variable genes in *P. berghei* females, 73 in males, 27 in trophozoites and four in schizonts. Twenty variable genes were shared between males and females. One variable gene from trophozoites and schizonts was shared: an L1 type *pir* gene (PBANKA_0943900). Ten were shared between females and trophozoites, none between females and schizonts, four between males and trophozoites, none between males and schizonts. To examine functional classes enriched amongst variable genes we used topGO with the weight01 algorithm, the Fisher statistic, node size = 5 and False Discovery Rate >= 0.05 ^48^. Gene ontology terms for *P. berghei* and *P. falciparum* genes were extracted from GeneDB EMBL files. Multigene families in *P. berghei* do not have associated Gene Ontology (GO) terms and so we used *ad hoc* hypergeometric tests to look at their enrichment. We found that *pir* genes were enriched amongst variable genes in trophozoites (2 of 135, hypergeometric test, *p*=0.04), schizonts (1 of 135, hypergeometric test *p*=0.005) and males (5 of 135, hypergeometric test, *p*=0.02), but there were none in females. Those in asexual stages (trophozoites and schizonts) were from the L1 subfamily, but those in males were from the S1 and S4 subfamilies.

Many more variable genes were present in *P. falciparum* females and so we reduced the FDR cut off to 0.01 to improve specificity. We found 448 variable genes in *P. falciparum* females. We were not able to analyse variability in *P. falciparum* males because we found only five of them. Amongst several enriched GO terms were *modulation by symbiont of host erythrocyte* and c*ytoadherence to microvasculature, mediated by symbiont protein*. These terms refer to 14 *var* genes found amongst the variable genes.

### Identifying putative functional *var* transcripts

It is known that antisense transcripts are expressed from a bidirectional promoter in the intron of each *var* gene ^17^. Our protocol does not preserve information about which strand is transcribed. Therefore finding that reads map to either exon of a *var* gene does not provide evidence that it is functionally expressed. In fact it may indicate that the gene is silenced. In order to identify sense transcripts, we looked for reads mapping over the single intron of each *var* gene. These reads that include both exons must originate from transcripts including the intron, and thus indicate that the gene was transcribed on the sense strand, not from antisense transcripts beginning within the intron. Initially we identified reads from the HISAT2 mappings which overlapped annotated *var* genes using *bedtools intersect*. From the resulting BAM file we selected those reads which included an *N* in the CIGAR string, indicating a split read. We then looked for which *var* gene each read overlapped and whether it was split exactly over the intron. We called expression for a *var* gene where there were at least two reads mapping over the intron.

## Code availability

Perl, R and C++ code for various analyses is available on request.

## Data Availability

The single-cell RNA-seq reads are available from the European Nucleotide Archive (accession ERP021229) and ArrayExpress (accession E-ERAD-611). Read counts and metadata are also presented in Supplementary Data 5 and 6.

## Acknowledgements

The Wellcome Trust Sanger Institute is funded by the Wellcome Trust (grant WT098051). CJRI was supported by a Sir Henry Dale Fellowship, jointly funded by the Wellcome Trust and the Royal Society (Grant Number 101239/Z/13/Z). MKNL is supported by an MRC Career Development Award (G1100339). We thank Chris Newbold and John Marioni for comments that improved the manuscript. The authors would like to thank the staff of the Illumina Bespoke Sequencing and Core Cytometry teams at the Wellcome Trust Sanger Institute for their contribution.

The authors declare no conflict of interest.

## Contributions

AJR, AMT, HMB and MKNL conceived of and designed the study. AMT and HMB carried out the sorting and library preparation protocols. AMT and ARG performed the parasitological experiments. AJR, AMT and HMB designed the sequencing experiments. AJR and AMT analysed the data. CJRI performed the Fourier transform analysis. MJS coordinated sequencing experiments. OB supervised ARG. AJR, AMT, MB & MKNL wrote the manuscript. All authors read and critically revised the manuscript.

## Supplementary Material

Supplementary Figure 1. Analysis of biases in *Plasmodium berghei* mixed blood stage single-cell RNA-seq libraries.

Supplementary Figure 2. Dual sorting of two parasite species shows minimal contaminating RNA

Supplementary Figure 3. Distribution of gene counts.

Supplementary Figure 4. The same subsets of transcripts show different patterns of expression around the end of the asexual cell cycle in conventional bulk RNA-seq data and pseudotime reconstructions of single cell RNAseq data.

Supplementary Figure 5. Pseudotime reconstruction of the late asexual trajectory of P. falciparum. Supplementary Figure 6. Analysis of the co-expression pattern of the ApiAP2 family of transcription factors (TFs).

Supplementary Figure 7. Principal Components Analysis and classification of P. falciparum gametocyte cells

Supplementary Table 1. Reagents permuted during optimisation of the single cell RNAseq protocol and stats of each treatment condition after sequencing.

Supplementary Data 1. Marker genes identifying *P. berghei* mixed stage k-means clusters Supplementary Data 2. Highly variable genes in *P. berghei* female gametocytes, male gametocytes, trophozoites, schizonts and *P. falciparum* female gametocytes.

Supplementary Data 3. GO terms enriched amongst highly variable *P. berghei and P. falciparum* genes

Supplementary Data 4. *Pir* gene expression in different cell types Supplementary Data 5. Samples sequenced in this study

Supplementary Data 6. Gene count tables for the three large datasets included in the study

## Supplementary Figures

**Supplementary Figure 1.**
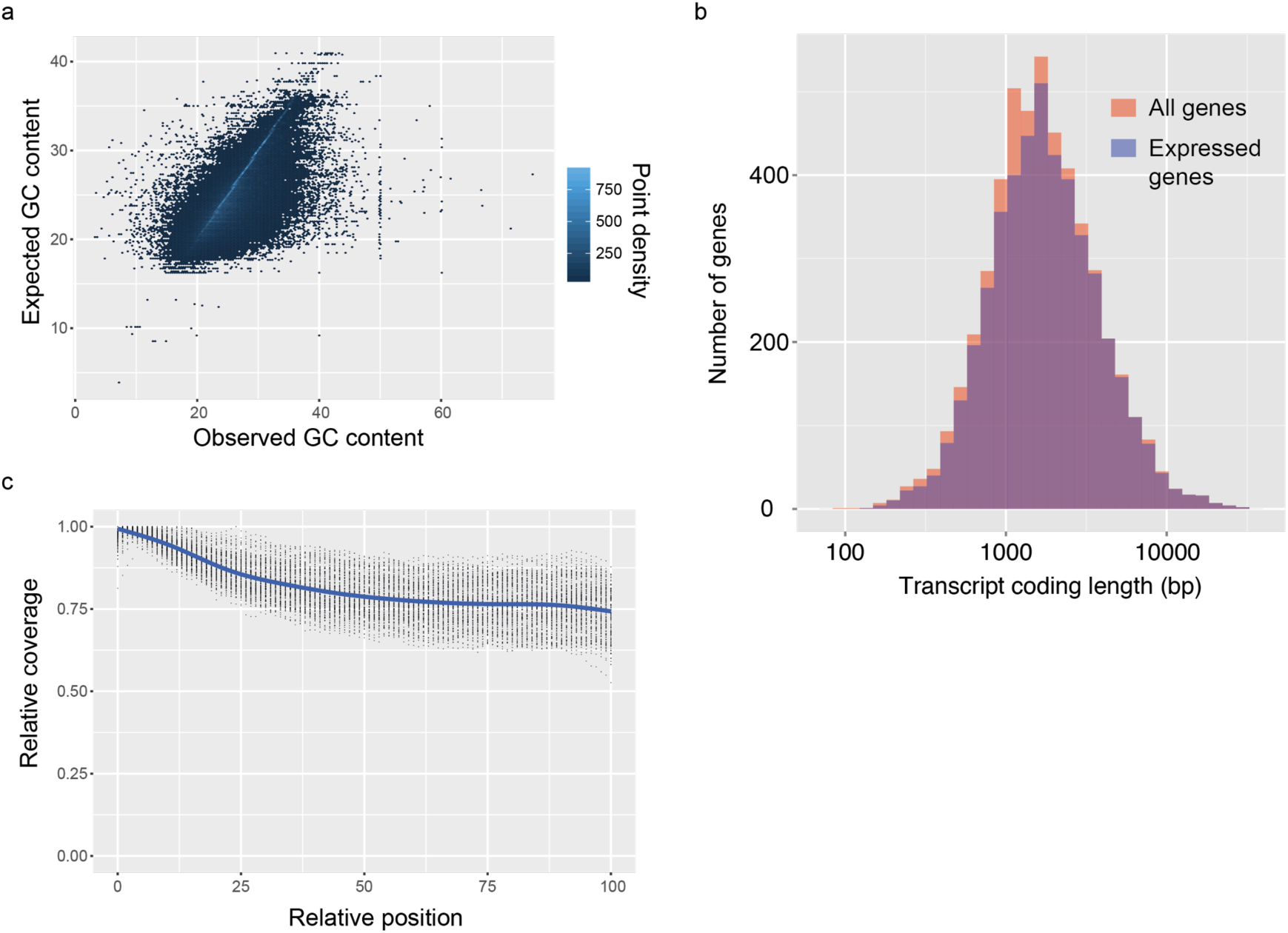
Analysis of biases in *Plasmodium* berghei mixed blood stage single-cell RNA-seq libraries. (a) The GC content of transcript fragments agreed well with the GC content of genes. There was no apparent over-or under-representation of GC rich regions. (b) Expressed genes (those with at least 10 reads in at least 5 cells) were representative of average gene length, suggesting that although the reverse transcriptase might not copy the whole of long transcripts, fragments of long genes are still detected. (c) Sequencing library preparation often introduces end bias, where either the 5’ or 3’ end of transcripts tend to be better covered. Our protocol introduced a small 5’-bias, which could be attributable to the reverse transcription sometimes initiating within transcripts in internal polyA regions, rather than in the 3’ poly-A tail.

**Supplementary Figure 2.**
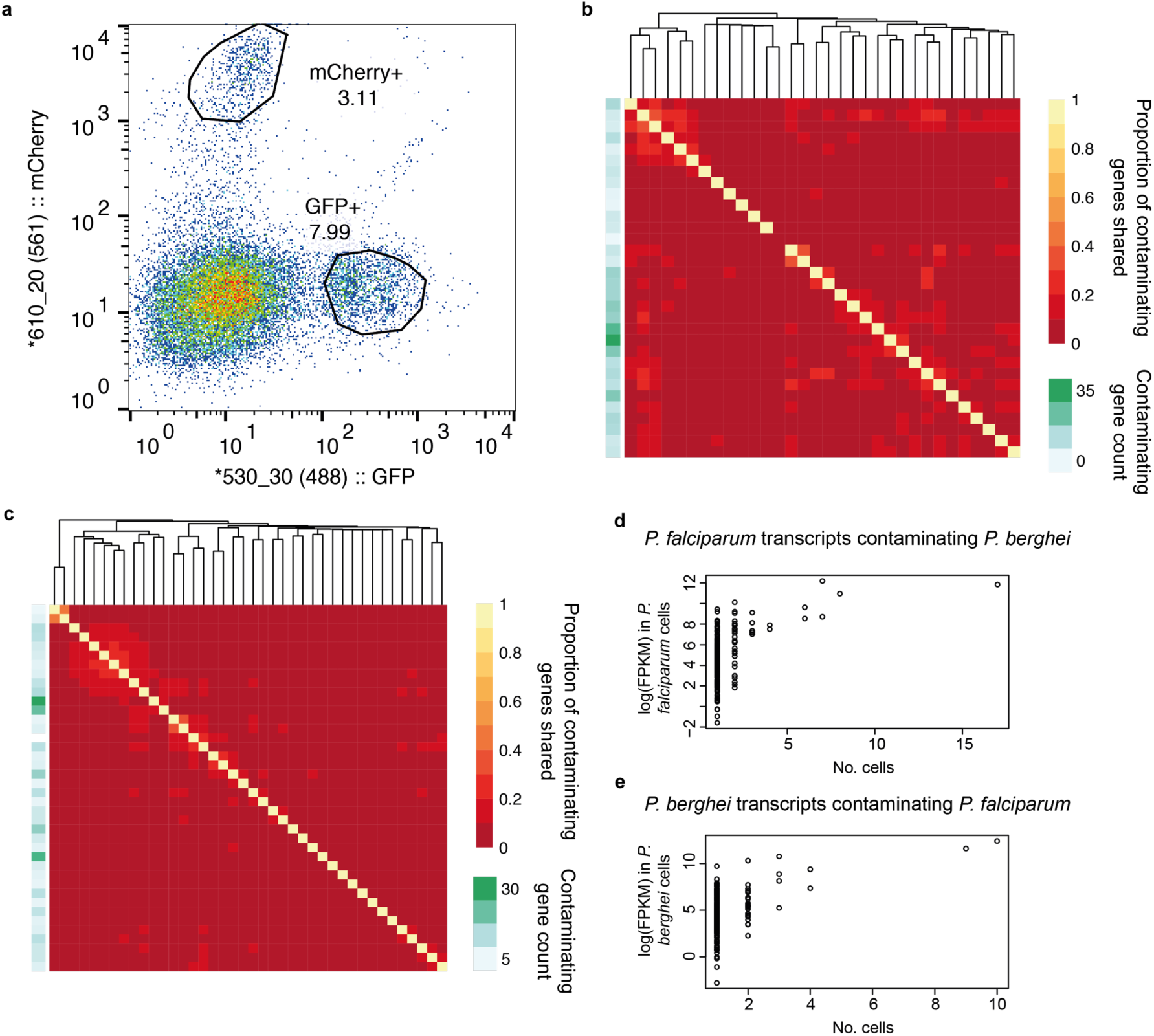
Dual sorting of two parasite species shows minimal contaminating RNA. (a) Purified asexual late blood stage of GFP *P. falciparum* and mCherry *P. berghei* were mixed at a 1:1 ratio, inactivated in RNALater, and sorted individually by flow cytometry, gated on respective fluorescent channels. The proportion of shared contaminating transcripts between pairs of cells was low for *P. falciparum* transcripts in *P. berghei* cells (b) and even lower for *P. berghei* transcripts in *P. falciparum* cells (c). Overall there were 273 unique *P. falciparum* transcripts contaminating the *P. berghei* transcriptomes, although no cell had more than 37 contaminating transcripts and no pair of cells shared more than 17. There was a total of 258 *P. berghei* transcripts contaminating *P. falciparum* transcriptomes, although no cell had more than 32 of these and no pair of cells shared more than 10. The data suggest that *P. falciparum* schizonts cause more contamination than *P. berghei* schizonts. More commonly occurring, contaminating transcripts are more highly expressed in their cells of origin for both *P. falciparum* transcripts contaminating *P. berghei* cells (d) and *P. berghei* transcripts contaminating *P. falciparum* cells (e). This suggests that contaminants reflect the observed pool of transcripts.

**Supplementary Figure 3.**
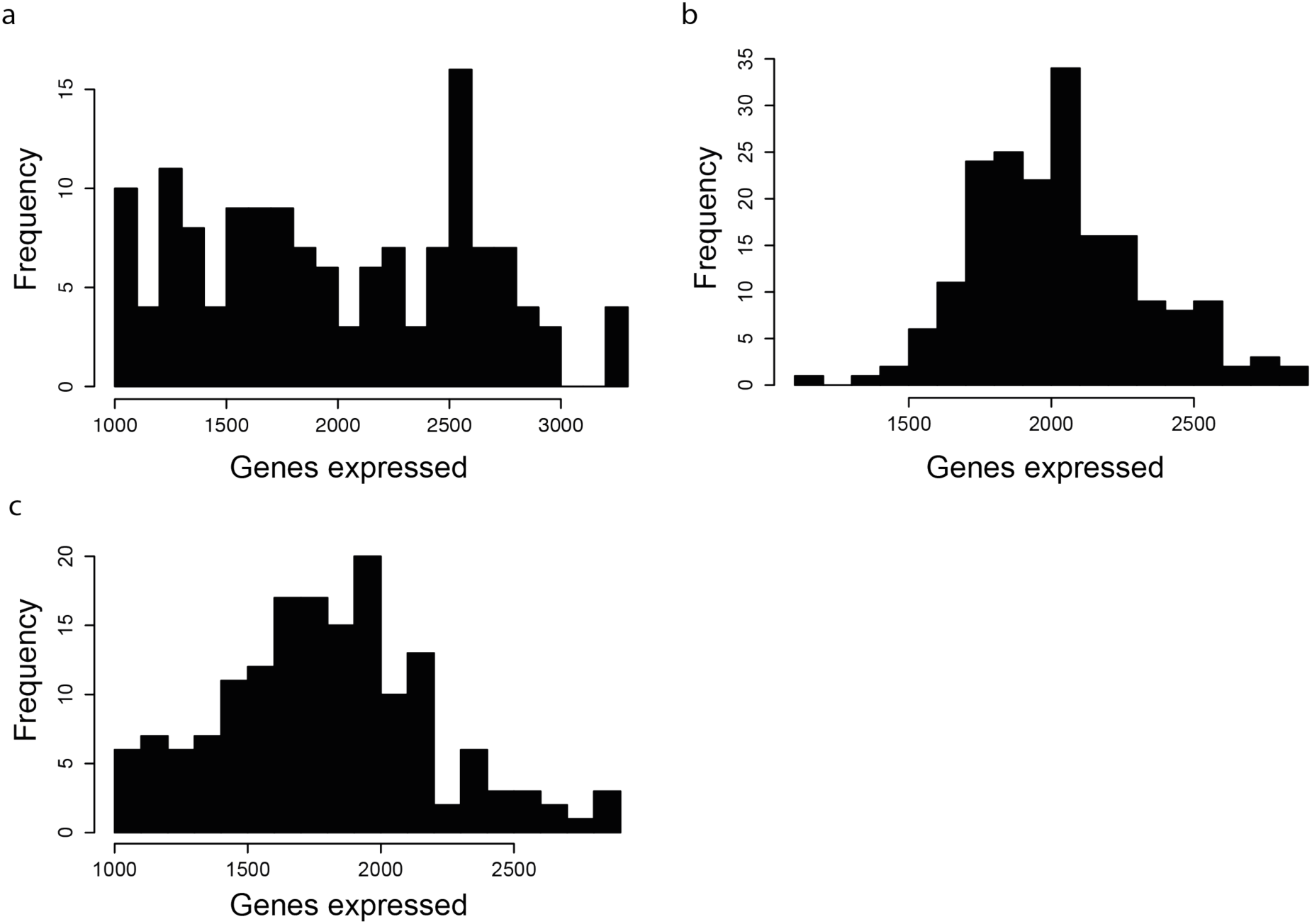
Distribution of gene counts. Histograms of expressed gene number after filtering for (a) 144 *P. berghei* mixed blood stage cells, (b) 191 *P. falciparum* gametocytes and (c) 161 *P. falciparum* asexual cells. The greater distribution of gene counts in *P. berghei* is due to the greater variety of cell types in this dataset. Female gametocytes for instance, consistently had a greater number of genes expressed.

**Supplementary Figure 4.**
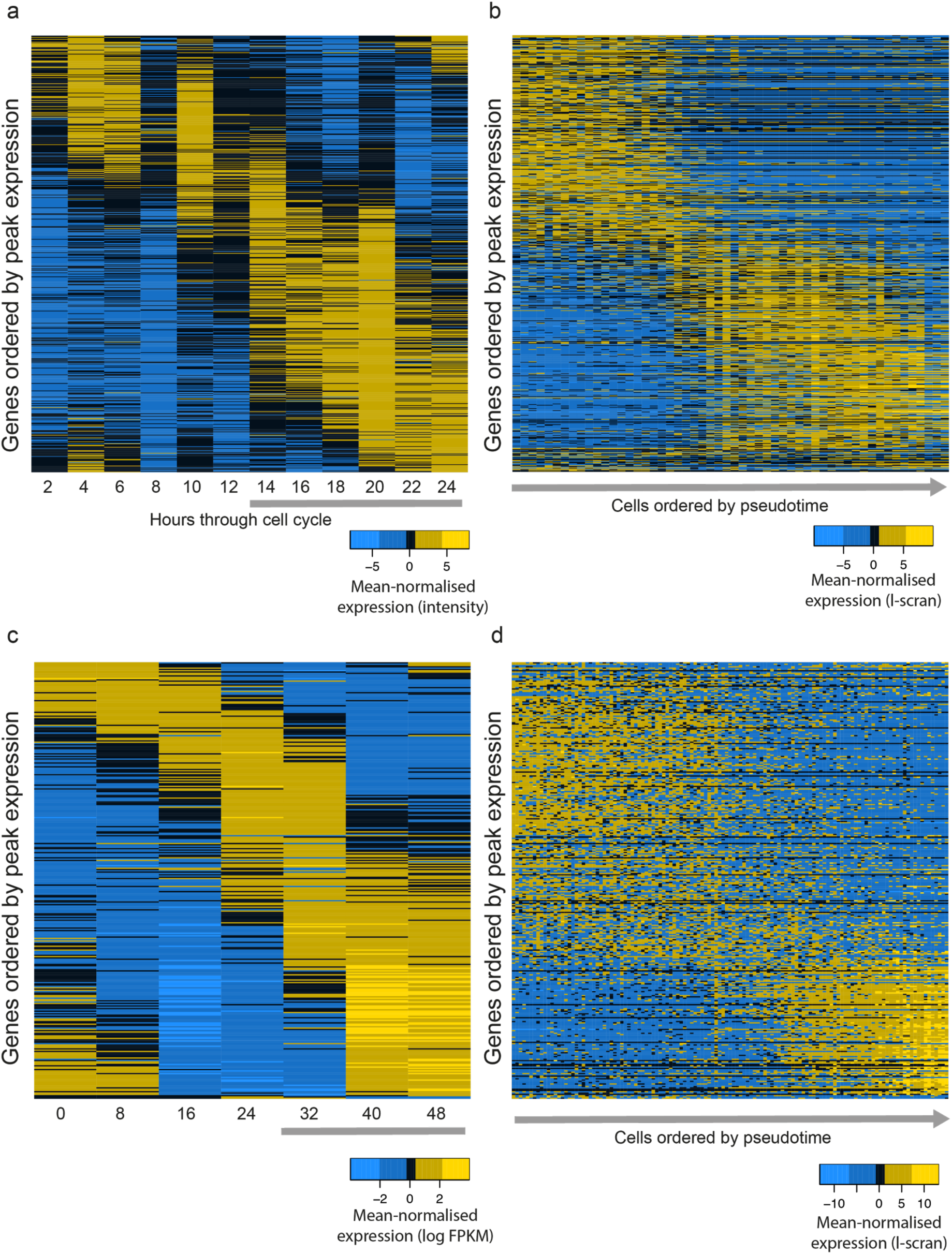
The same subsets of transcripts show different patterns of expression around the end of the asexual cell cycle in conventional bulk RNA-seq data and pseudotime reconstructions of single cell RNAseq data. A shared set of 651 genes identified as following a sigmoidal expression pattern through the intraerythrocytic developmental cycle (see Methods) are shown in both bulk transcriptome data ^11^ (**a**) and single cell data ordered by pseudotime (**b**) for *P. berghei*. A much more dramatic shift in gene expression is observed in the single-cell transcriptome data. A similar pattern is observed between *P. falciparum* bulk^24^ (**c**) and single cell (**d**) RNA-seq. In panels b and d, gene expression patterns are mean-normalised l-scran values. Only late stage parasites (grey bars in bulk reference datasets) are expected to be present in the single cell datasets.

**Supplementary Figure 5.**
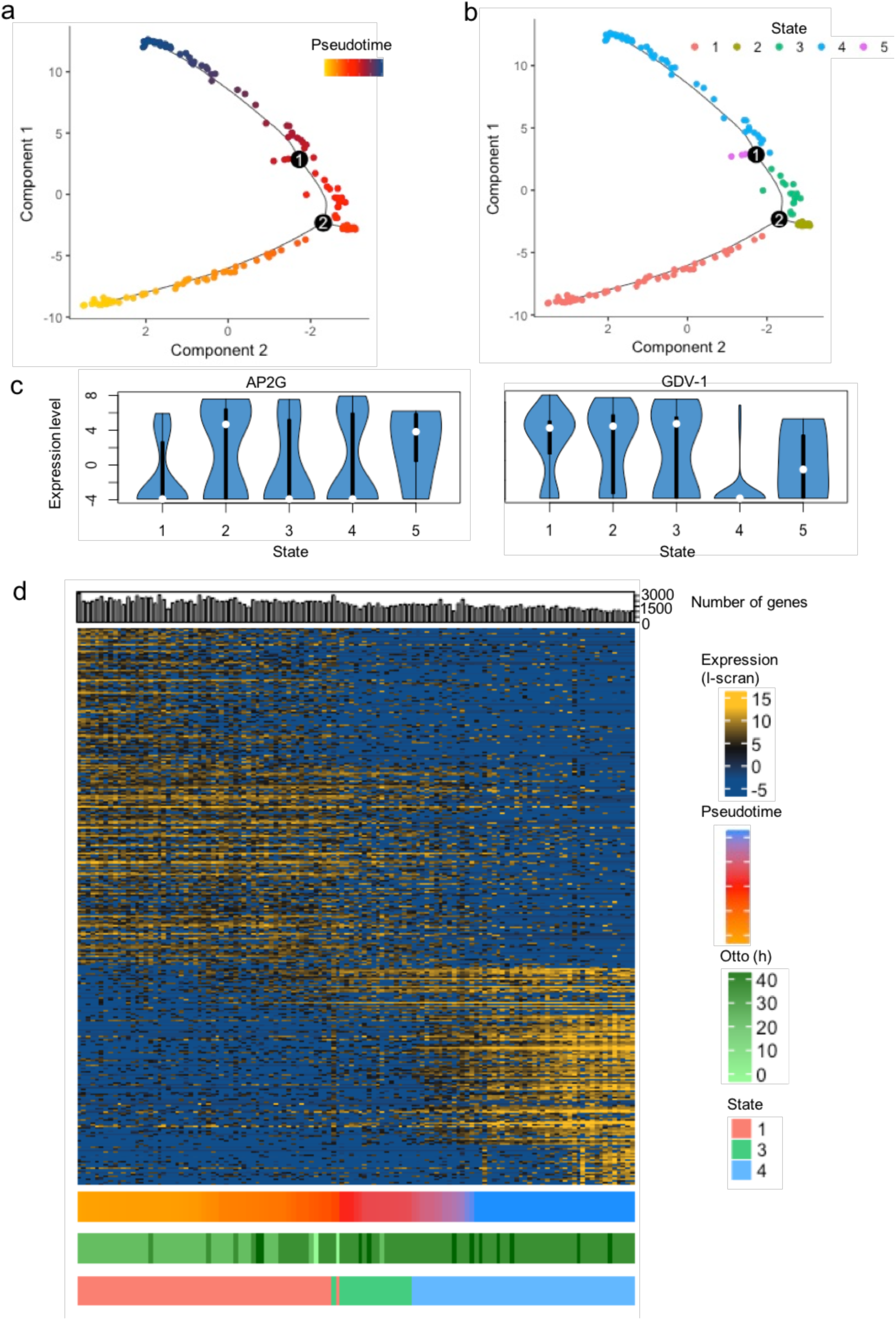
Pseudotime reconstruction of the late asexual trajectory of *P. falciparum*. PCA of 155 *P. falciparum* cells colored by pseudotime (**a**) or Monocle state (**b**); identified trajectory branches are displayed as circled number 1 and 2. (**c**) Expression of sexual commitment markers *ap2-g* (PF3D7_1222600) and *gdv-1* (PF3D7_0935400) in cells of different states. (**d**) Differentially expressed genes were plotted along pseudotime for cells in the main trajectory (States 1, 3 and 4). The number of genes per cell is displayed on top of the heat map, whilst the pseudotime, the maturation prediction ^24^ and the cell state are displayed on the side of the heatmap. The transition between trophozoites and schizonts is associated with a hard transcriptional shift, as seen for *P. berghei* (Fig. 2d).

**Supplementary Figure 6.**
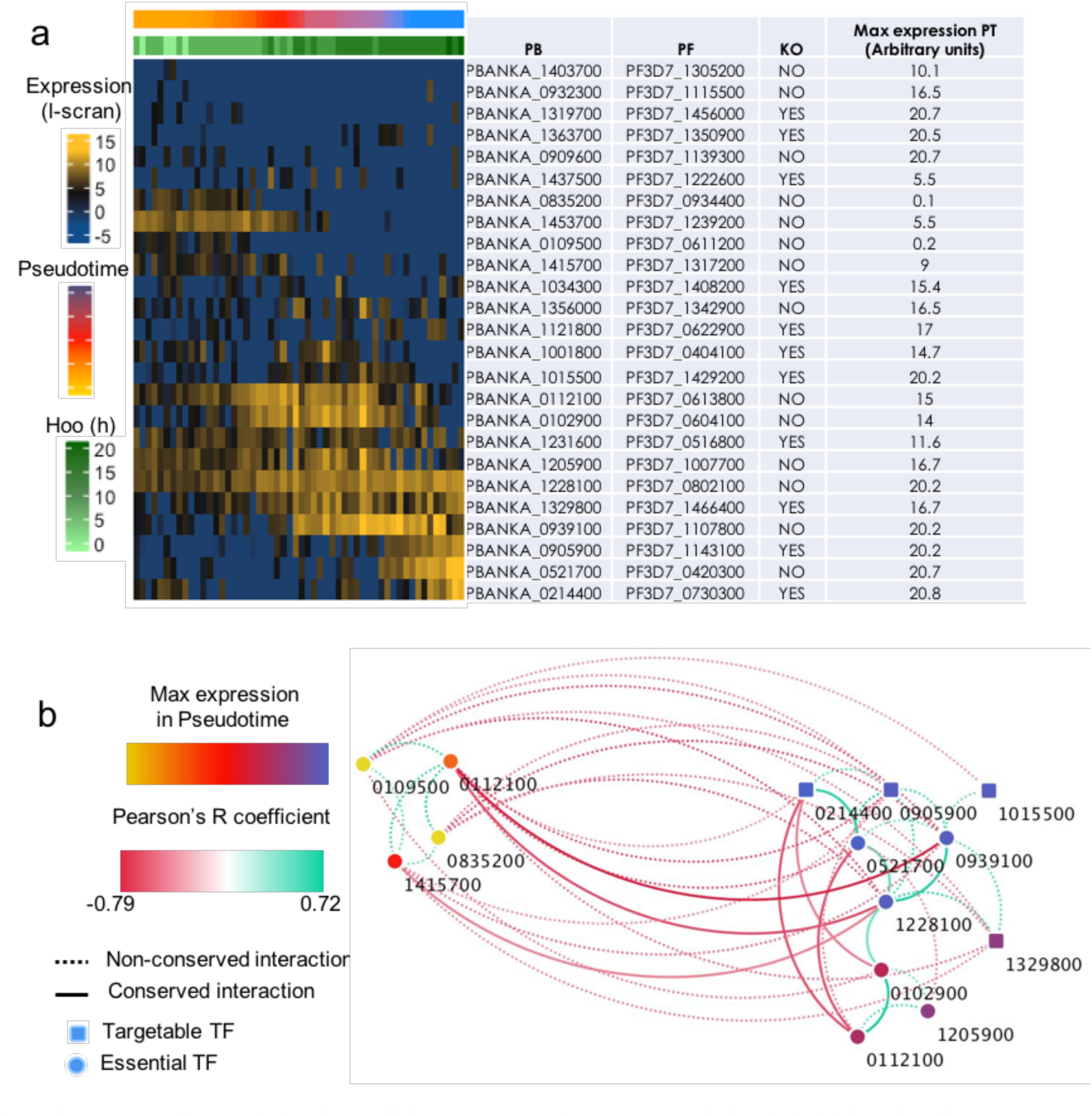
Analysis of the co-expression pattern of the ApiAP2 family of transcription factors (TFs). (a) Expression of *P. berghei* ApiAP2 genes in pseudotime. The *P. falciparum* ortholog is indicated (PF), as well as its established or custom short name (Pb), it’s peak pseudotime expression (Max expression PT) and the ability to disrupt the TF as observed in a recent genetic screen (KO) ^49^. (b) A network analysis was conducted using significant positive and negative interactions (*p* < 0.05 by Pearson’s correlation) of 25 different TFs and weighted according to their Pearson correlation coefficient. A similar analysis in *P. falciparum* revealed that some of these co-expression interactions are conserved within the genus (solid lines). Maximum TF expression along pseudotime appears to be important to the structure of the network, strongly suggesting a coordinated regulation cascade of different members of the family during the trophozoite to schizont transition.

**Supplementary Figure 7.**
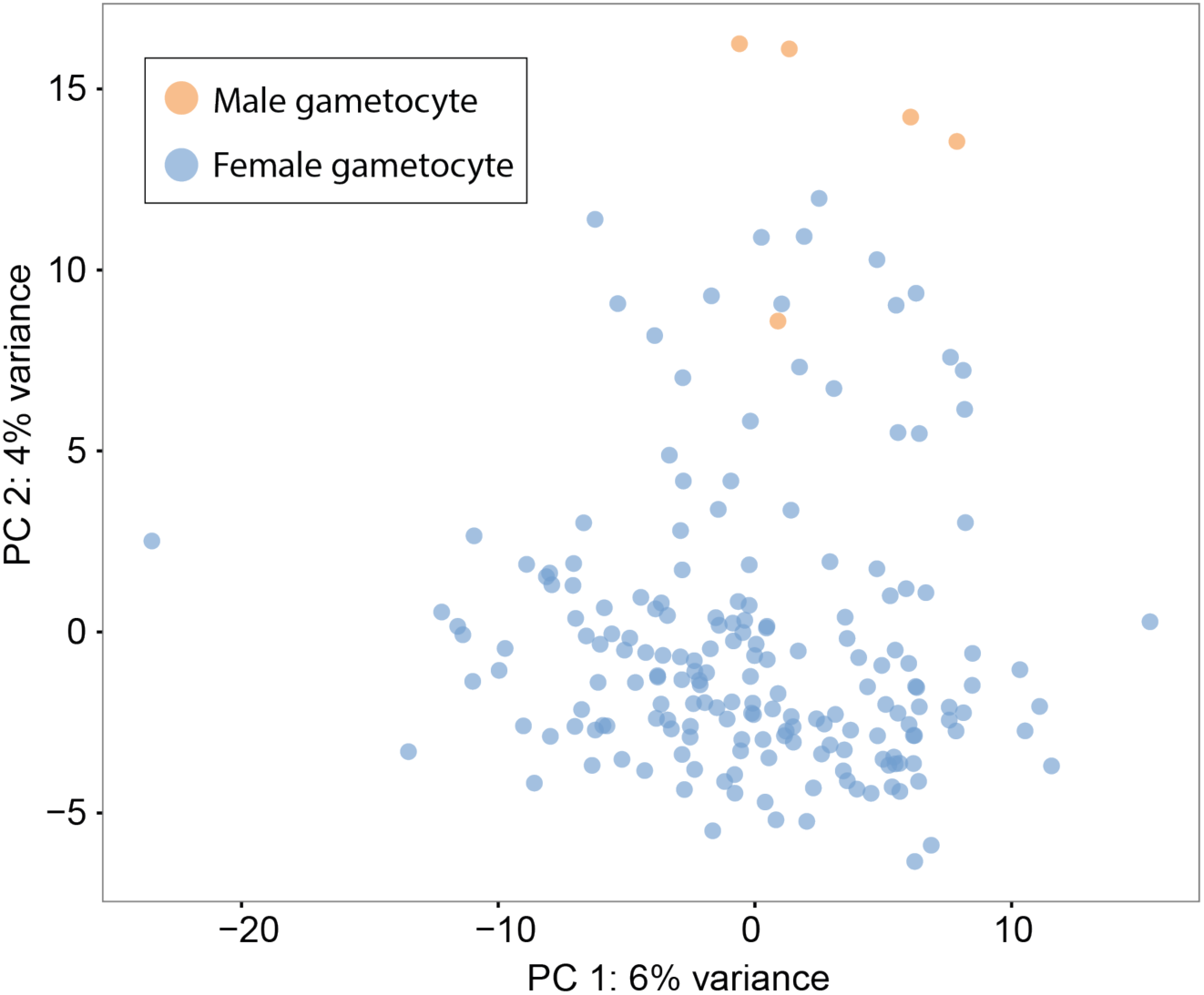
Principal Components Analysis and classification of *P. falciparum* gametocyte cells. A combination of Principal Components Analysis (PCA), k-means clustering and comparison to bulk RNA-seq datasets was used to classify 191 high quality *P. falciparum* gametocytes. A consensus of clustering and comparison to bulk RNA-seq allowed us to distinguish male gametocytes and female gametocytes.

### Supplementary Tables/Data

**Supplementary Table 1.** Reagents permuted during optimisation of the single cell RNAseq protocol and stats of each treatment condition after sequencing.

| Conditions tested | Protocol | SSII, V30, 30 cycles | SSII, T30, 30 cycles | SmSc, T30, 30 cycles | SSII, T30, 25 cycles | SmSc, T30, 25 cycles | SmSc, T30, 25 cycles | SmSc, T30, 25 cycles | SmSc, T30, 25 cycles |
| --- | --- | --- | --- | --- | --- | --- | --- | --- | --- |
|  | Cells | Sexual | Asexual | Asexual | Asexual | Asexual | Sexual | Mixed blood | Asexual |
|  | Species | Pf | Pf | Pf | Pf | Pf | Pf | Pb | Pf |
| Lysis buffer volume | 2 µl | ✓ |  |  |  |  |  |  |  |
|  | 4 µl |  | ✓ | ✓ | ✓ | ✓ | ✓ | ✓ | ✓ |
| Oligo Dt (IDT) | Anchored 30 bp | ✓ |  |  |  |  |  |  |  |
|  | Non-Anchored 30 bp |  | ✓ | ✓ | ✓ | ✓ | ✓ | ✓ | ✓ |
| Reverse transcriptase | Superscript II (Life Technologies) 10U | ✓ | ✓ |  | ✓ |  |  |  |  |
|  | Smartscribe (Clontech) 5U |  |  | ✓ |  | ✓ | ✓ | ✓ | ✓ |
| Cycle number | 25 |  |  |  | ✓ | ✓ | ✓ | ✓ | ✓ |
|  | 30 | ✓ | ✓ | ✓ |  |  |  |  |  |
| Sequencing machine | HiSeq |  |  |  |  |  | ✓ | ✓ | ✓ |
|  | MiSeq | ✓ | ✓ | ✓ | ✓ | ✓ |  |  |  |
| Sequencing results summary | % rRNA | 5.7 | 33.5 | 36.2 | 6.4 | 18.4 | 17.8 | 16.7 | 34.8 |
|  | % coding genes | 4.4 | 11.3 | 39.3 | 10.5 | 33 | 51.7 | 49 | 40.5 |
|  | % other | 90 | 55.2 | 24.4 | 83.1 | 48.6 | 30.5 | 34.2 | 24.6 |
|  | Median genes detected for 50k reads | 25 | 84 | 145 | 174 | 181 | 502.5 | NA | NA |
|  | Total cells | 5 | 6 | 6 | 6 | 6 | 237 | 182 | 174 |
|  | Cells passing filters | NA | NA | NA | NA | NA | 191 | 144 | 161 |
|  | Median gene count | NA | NA | NA | NA | NA | 2011 | 1922.5 | 1793 |

Different combinations of the protocol were tested by sequencing. Initial trials were performed with 2 µl of lysis buffer, this was increased to 4µl to augment capture efficiency. Permutations of the protocol that were tested were a terminal anchoring base (A,G,C; **V**) or not (**T**), 2 reverse transcriptase enzymes (Smartscribe (SmSc); Superscript II (SII)) and 25 or 30 cycles of preamplification. Both sexual and asexual cells of *P. berghei* and *P. falciparum* were tested. For each sequenced dataset, we calculated the mean percentages of rRNA, mRNA and other reads across the cells. For some samples we also downsampled the data to 50,000 reads per cell to allow comparison of the number of genes detected. This was done to determine differences in the complexity of each library. For the three larger datasets produced (*P. falciparum* gametocytes, *P. berghei* mixed blood stages, and *P. falciparum* asexual stages), we provide the numbers of pre-and post-filtered cells and median number of genes in those filtered cells.

**Supplementary Data 1. Marker genes identifying *P. berghei* mixed stage k-means clusters** Marker genes are those for which expression is reliably associated with a particular cluster. They were identified for clusters in the k = 3 means clustering of *P. berghei* mixed blood stages using the *markers* function of the SC3 package. Cluster 1 corresponds to asexuals, cluster 2 to males and cluster 3 to females. The majority consensus is the cell type most common in that cluster. However, the only alternative cells in these clusters are the small number of outliers. AUROC is the Area Under the Receiver Operating Characteristics curve and is a measure of the reliability of the marker. We used an AUROC cut off of 0.85 and a p-value cut off of 0.01.

*See Excel file “Supplementary Data 1.xls”*

**Supplementary Data 2. Highly variable genes in *P. berghei* female gametocytes, male gametocytes, trophozoites, schizonts and *P. falciparum* female gametocytes**. The *p* and *q* (corrected) values given for each gene are those determined using M3Drop.

See Excel file “Supplementary Data 2.xls”

**Supplementary Data 3. Gene Ontology terms enriched amongst highly variable *P. berghei and P. falciparum* genes.**

Enriched GO terms were determined using topGO. The total terms are those in the whole set of transcripts and multiple-hypothesis testing corrected significance values (q values) are shown.

See Excel file “Supplementary data 3.xls”

**Supplementary Data 4. Pir gene expression in different P. berghei cell types**

*Pir* gene expression values for *P. berghei* mixed blood stages are shown, relating to Figure 3a.

Expression is measured by length normalised scran, or l-scran values.

See Excel file “Supplementary Data 4.xls”

**Supplementary Data 5. Samples sequenced in this study**

A full list of the samples related to this study is presented, linking the data we present to sequence data identifiers used by the European Nucleotide Archive (sanger_sample_id). Type: SC = single cell, Hcell = 100 cells, NoCell = no cell control. Run = internal Illumina run number. Lane = Illumina lane number. Tag = Internal Nextera barcode number. is_control = is the sample a control sample? Pass_filter = did the sample pass our filtering criteria. consensus = consensus annotation for that cell. For P. berghei mixed blood stages: sc3_k4 = cluster identifier for k means clustering (k=4), sc3_k3 = cluster identifier for k means clustering (k=3), hoo = best matching sample from Hoo et al.^11^ bulk microarray dataset of cell cycle, otto = best matching sample from Otto et al.^12^ bulk RNA-seq dataset of life cycle stages. For P. falciparum: sc3_female_k3 = cluster identifier for k means clustering of female-only samples (k=3), lasonder = best matching samples from ^42^ bulk RNA-seq dataset of male and female parasites, young = best matching samples from Young et al. bulk microarray dataset ^44^ of gametocyte development. For P. falciparum asexual stages, lopez = best matching sample from Lopez-Barragnan et al.^45^, otto = best matching sample from Otto et al.^24^ and pseudotime_state = pseudotime state, where states 2 and 5 were filtered out.

See Excel file “Supplementary Data 5.xls’

**Supplementary Data 6. Gene counts for the datasets included in the study**

Gene counts for all cells (included those we failed) for the P. berghei mixed blood stages, theP. falciparum gametocytes, and the P. falciparum asexuals are presented here. These data, along with the corresponding metadata in Supplementary Data 5 can be used to replicate and/or extend our analysis using, for instance, the scater package ^50^.

See Excel file “Supplementary Data 6.xls’

